# Hepatocyte cholesterol content modulates glucagon receptor signalling

**DOI:** 10.1101/2021.10.31.466084

**Authors:** Emma Rose McGlone, T. Bertie Ansell, Cecilia Dunsterville, Wanling Song, David Carling, Alejandra Tomas, Stephen R Bloom, Mark S. P. Sansom, Tricia Tan, Ben Jones

## Abstract

Glucagon decreases liver fat, and non-alcoholic fatty liver disease (NAFLD) is associated with hepatic glucagon resistance. Increasingly it is recognised that the function of G protein-coupled receptors can be regulated by their local plasma membrane lipid environment. The aim of this study was to evaluate the effects of experimentally modulating hepatocyte cholesterol content on the function of the glucagon receptor (GCGR). We found that glucagon-mediated cAMP production is inversely proportional to cholesterol content of human hepatoma and primary mouse hepatocytes after treatment with cholesterol-depleting and loading agents, with ligand internalisation showing the opposite trend. Mice fed a high cholesterol diet had increased hepatic cholesterol and a blunted hyperglycaemic response to glucagon, both of which were partially reversed by simvastatin. Molecular dynamics simulations identified potential membrane-exposed cholesterol binding sites on the GCGR. Overall, our data suggest that increased hepatocyte membrane cholesterol could directly contribute to glucagon resistance in NAFLD.

## Introduction

Glucagon is a key hormone in the control of hepatic metabolism: as well as increasing glucose production to counteract hypoglycaemia, it reduces liver fat by decreasing *de novo* lipogenesis and increasing fatty acid oxidation (Longuet et al., 2008; Penhos et al., 1966). Patients with type 2 diabetes mellitus (T2DM) and/or non-alcoholic fatty liver disease (NAFLD) exhibit hepatic glucagon resistance (Demant et al., 2018; Suppli et al., 2020), which in turn contributes to worsening of steatosis by blocking the beneficial effects of glucagon on hepatic lipid metabolism. To date, the mechanism behind this phenomenon is incompletely understood (Janah et al., 2019). Hepatic cholesterol accumulation is a feature of NAFLD and, interestingly, the degree of both glucagon resistance (Gar et al., 2021) and hepatic cholesterol accumulation (Min et al., 2012; Puri et al., 2007) are correlated with histological severity of the disease.

The glucagon receptor (GCGR) is a prototypical class B G protein-coupled receptor (GPCR). It is increasingly appreciated that the functions of cell surface GPCRs are heavily modulated by other membrane components (Thal et al., 2018) including lipids (Reiter et al., 2017). This can occur either through direct lipid-receptor interactions (Duncan et al., 2020b), or indirectly, e.g. by altering the properties of the membrane bilayer (Mahmood et al., 2013). Cholesterol is known to alter the conformation of some GPCRs by directly binding to allosteric sites (Casiraghi et al., 2016; Manna et al., 2016); it is also a key structural component of the plasma membrane that controls the distribution and spatial coupling between surface receptors and intracellular mediators (Simons and Ikonen, 2000). We have recently demonstrated that cholesterol depletion affects signalling and endomembrane trafficking of the closely related glucagon-like peptide 1 receptor (GLP-1R) (Buenaventura et al., 2019). To date, the relevance of cellular cholesterol to GCGR signalling has not been explored experimentally.

The aim of this study was to investigate whether hepatocyte cholesterol content affects GCGR signalling and its downstream physiological correlates. Here we demonstrate that enrichment of cholesterol *in vitro* and *in vivo* decreases glucagon responsiveness. Cholesterol depletion has the opposite effect. Molecular dynamics simulations identify likely cholesterol binding sites on the GCGR which may allosterically regulate its function. Our results indicate that hepatocyte cholesterol content contributes to hepatic glucagon sensitivity and pinpoint a likely molecular basis for this phenomenon. Increased cellular cholesterol, by directly interacting with the hepatic GCGR, could contribute to glucagon resistance in patients with NAFLD.

## Results

### Manipulating cellular cholesterol content has divergent effects on GCGR-mediated signalling and ligand uptake

We first examined the impact of pharmacological modulation of cellular cholesterol levels on GCGR function. As hepatoma cells lack the high levels of GCGR expression typical of primary hepatocytes, we used Huh7 cells with exogenous GCGR expression for these studies (McGlone et al., 2021). cAMP signalling responses to glucagon stimulation in Huh7-GCGR cells treated with different concentrations of cholesterol-free methyl-β-cyclodextrin (MβCD), which rapidly extracts cholesterol from cellular membranes, showed an increase in potency and, at the highest MβCD concentration (10 mM), a trend (p=0.06) towards increased maximum response (Figure 1A; Table 1). Conversely, incubating with cholesterol-saturated MβCD to increase membrane cholesterol levels caused a reduction in cAMP signalling (Figure 1A; Table 1). Pre-treatment of Huh7-GCGR cells with simvastatin to inhibit cellular cholesterol synthesis also increased potency for cAMP in response to glucagon (Figure 1B; Table 1). This effect was reversed by concurrent pre-treatment with mevalonate to replenish the product of the statin-inhibited 3-hydroxy-3-methyl-glutaryl-coenzyme A (HMG-CoA) reductase reaction, and by acute restoration of membrane cholesterol levels using cholesterol-saturated MβCD. When we compared the relative cellular cholesterol content obtained in these assays against corresponding cAMP responses, a robust inverse correlation was seen (R^2^ =0.78, p=0.0003; Figure 1C).

**Table 1:** cAMP responses to glucagon and FITC-GCG uptake in Huh7-GCGR cells following cellular cholesterol manipulation. Parameter estimates ± SEM from responses depicted in Figure 1. For uptake assays, E_max_ and EC_50_ are derived from pooled data before statistical analysis. Mean ± SEM; *n*=4 or 5. Uptake at a single concentration of FITC-GCG (200 nM) is presented because E_max_ and EC_50_ could not be derived from the curves for overnight treatments. Treatment effects were analysed using matched 1-way ANOVA and compared to vehicle with Dunnett’s multiple comparison test, or by paired t-test where there are only 2 groups. “n.d.” indicates not done. *p<0.05; **p<0.01; ***p<0.001.

| Acute treatments |  |  |  |  |  |  |  |
| --- | --- | --- | --- | --- | --- | --- | --- |
|  |  | Vehicle | MβCD (1 mM) | MβCD (3 mM) | MβCD (10 mM) | Cholesterol |  |
| cAMP (Fig 1A) | Log EC<sub>50</sub> (M) | -10.5 ± 0.0 | -10.7 ± 0.0 * | -10.8 ± 0.1 * | -10.6 ± 0.1 | -10.2 ± 0.0 * |  |
|  | E<sub>max</sub> (nM cAMP) | 14.9 ± 2.7 | 15.0 ± 1.2 | 16.1 ± 0.5 | 21.9 ± 2.4 | 14.8 ± 1.6 |  |
| Gα<sub>s</sub> (Fig 1D) | Log EC<sub>50</sub> (M) | -10.4 ± 0.1 | n.d. | -10.5 ± 0.1 | n.d. | -10.0 ± 0.1 ** |  |
|  | E<sub>max</sub> (nM cAMP) | 15.4 ± 1.0 | n.d. | 31.3 ± 3.0 *** | n.d. | 19.6 ± 2.0 |  |
| Gα<sub>i</sub> (Fig 1D) | Log EC<sub>50</sub> (M) | -9.1 ± 0.1 | n.d. | -9.4 ± 0.2 | n.d. | -9.2 ± 0.3 |  |
|  | E<sub>max</sub> (nM cAMP) | 13.4 ± 1.0 | n.d. | 26.1 ± 3.0 *** | n.d. | 16.6 ± 2.0 |  |
| GCG-FITC uptake (Fig 1G) | Log EC<sub>50</sub> (M) | -6.9 ± 0.2 | -6.8 ± 0.1 | -6.2 ± 0.2 ** | n.d. | -6.9 ± 0.1 |  |
|  | E<sub>max</sub> (RFU) | 89.4 ± 8.0 | 60.1 ± 3.0* | 56.5 ± 8.4 * | n.d. | 133.0 ± 8.0** |  |
|  | 200 nM (RFU) | 62.5 ± 6.3 | 38.8 ± 2.0 * | 24.6 ± 1.8 *** | n.d. | 87.8 ± 7.0 ** |  |
| Overnight treatments |  |  |  |  |  |  |  |
|  |  | SFM | SFM + Chol | SFM + Mev | Simva | Simva + Chol | Simva + Mev |
| cAMP (Fig 1B) | Log EC<sub>50</sub> (M) | -10.5 ± 0.1 | -10.0 ± 0.1 * | -10.3 ± 0.0 | -10.7 ± 0.1 * | -10.3 ± 0.1 | -10.5 ± 0.0 |
|  | E<sub>max</sub> (% FSK) | 61.4 ± 14.1 | 69.0 ± 18.6 | 62.8 ± 16.0 | 83.8 ± 24.3 | 82.5 ± 29.9 | 50.9 ± 14.9 |
| Gα<sub>s</sub> (Fig 1E) | Log EC<sub>50</sub> (M) | -10.2 ± 0.1 | n.d. | n.d. | -10.4 ± 0.1 *** | n.d. | n.d. |
|  | E<sub>max</sub> (% FSK) | 84.0 ± 11.5 | n.d. | n.d. | 98.6 ± 12.4 | n.d. | n.d. |
| Gα<sub>i</sub> (Fig 1E) | Log EC<sub>50</sub> (M) | -9.5 ± 0.1 | n.d. | n.d. | -9.5 ± 0.3 | n.d. | n.d. |
|  | E<sub>max</sub> (% FSK) | 73.7 ± 11.4 | n.d. | n.d. | 90.9 ± 14.0 | n.d. | n.d. |
| GCG-FITC uptake (Fig 1I) | 200 nM (RFU) | 78.5 ± 13.4 | 98.6 ± 17.7 | 85.7 ± 10.5 | 89.2 ± 12.8 | 85.1 ± 19.2 | 79.8 ± 6.9 |

**Figure 1:**
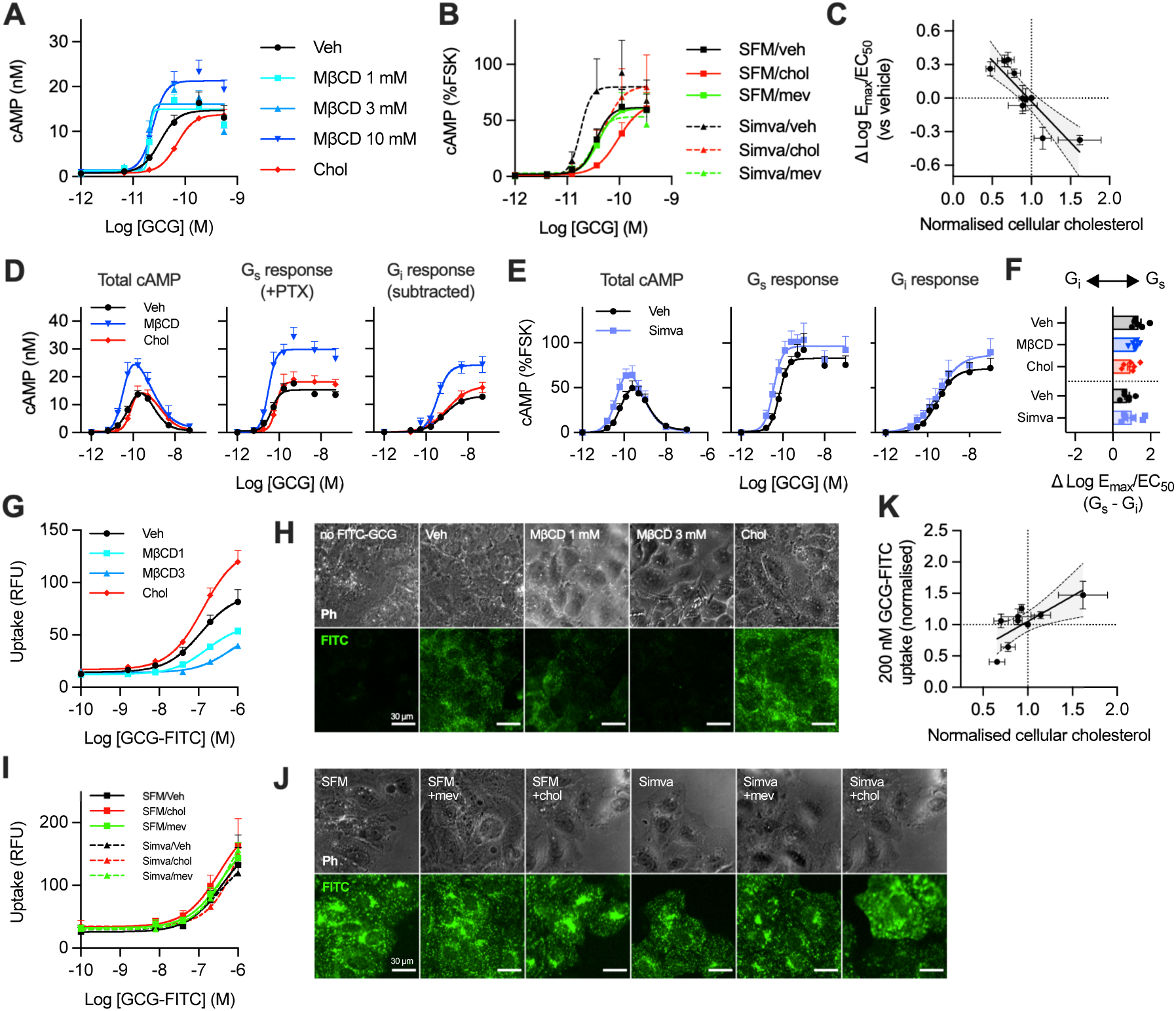
Increasing cellular cholesterol decreases agonist-stimulated cAMP production at GCGR whilst increasing agonist uptake in vitro. (**A**) cAMP concentration response curves in Huh7-GCGR cells pre-treated with cholesterol-deplete MβCD or cholesterol-saturated MβCD (Chol; 50 µg/ml), then stimulated with glucagon (GCG) n=4; or (**B**) pre-treated with simvastatin (Simva) or serum-free medium (SFM) overnight, with concurrent or subsequent treatment with or without mevalonate (mev; 50 µM) or cholesterol-saturated MβCD, then stimulated with glucagon. Results in (B) were normalised to forskolin responses (FSK; 10 µM), n=5. (**C**) Association between a combined measure of cAMP efficacy and potency (log-transformed E_max_/EC_50_) and cellular cholesterol for each of the treatments shown in (A) and (B), both normalised to vehicle control, with linear regression line ± 95% confidence intervals shown. (**D**) cAMP responses over a wider glucagon concentration range with or without pertussis toxin pre-treatment (PTX; 10 ng/ml) to block Gα_i_-mediated cAMP inhibition and reveal the Gα_s_-specific response, n=6. The effect of pre-treatment with MβCD or cholesterol is shown. (**E**) As for (D) but the effect of overnight pre-treatment with simvastatin, n=6. (**F**) Balance between Gα_s_ and Gα_i_-mediated cAMP effects from (D) and (E); all inter-group statistical comparisons non-significant. (**G**) The effect of pre-treatment with MβCD or cholesterol on FITC-GCG uptake in Huh7-GCGR cells, n=5; with representative phase contrast (Ph) and FITC-epifluorescence images shown in (**H**). Scale bar = 30 µm. (**I**) and (**J**) are similar to (G) and (H) but showing the effect of simvastatin ± mevalonate ± cholesterol, n=4, scale bar = 30 µm. (**K**) Association between FITC-GCG uptake at 200 nM and cellular cholesterol content for each of the treatments shown in (G) and (I), with linear regression line ± 95% confidence intervals shown. Data are shown as mean + or ± SEM.

GCGR shows a biphasic cAMP concentration-relationship due to superimposed effects of Gα_s_-mediated stimulation and Gα_i_-mediated inhibition of cAMP production (Grady et al., 1987). We investigated the effects of modifying cellular cholesterol content on the Gα_s_ and Gα_i_ components of the overall cAMP response by performing experiments with and without the Gα_i_ inhibitor pertussis toxin (PTX) (Yajima et al., 1978) (Figure 1D, 1E, Table 1). The Gα_i_ response required at least 10-fold higher glucagon concentrations than the Gα_s_ response, and the balance between each component was not significantly affected by acute cholesterol depletion or loading, or by sustained cholesterol reduction with simvastatin (Figure 1F).

GCGR ligand internalisation represents an additional downstream functional readout of GCGR activation. We examined this phenomenon in Huh7-GCGR cells following treatment with cholesterol lowering or enriching agents using fluorescein isothiocyanate-tagged glucagon (“FITC-GCG”), which closely mimics the pharmacology of native glucagon (McGlone *et al*., 2021). In contrast to the effects on cAMP signalling, we found an increase in FITC-GCG uptake after cholesterol loading, and a decrease after cholesterol depletion (Figure 1G, 1H, Table 1). In experiments designed to determine the effect of simvastatin pre-treatment, FITC-GCG uptake showed lower potency for all conditions, precluding accurate assessment of the effects of each treatment on FITC-GCG uptake EC_50_ (Figure 1I, 1J). This may have been a result of the prolonged serum starvation period typically used in simvastatin incubations, which itself leads to loss of membrane cholesterol due to the absence of serum lipoproteins. Across all assays, a positive correlation was observed between FITC-GCG uptake and cellular cholesterol

### Molecular dynamics simulations reveal potential GCGR-cholesterol binding sites

To explore the potential for direct interactions between the GCGR and cholesterol in the plasma membrane as an underlying mechanism for our pharmacological observations, we performed coarse-grained molecular dynamics (MD) simulations of GCGR interactions with bilayer lipids when in active and inactive states. A single receptor molecule was simulated whilst embedded in plasma membrane mimetic bilayers containing 25% cholesterol. The locations of predicted cholesterol binding sites with the four highest cholesterol residence times were the same for the two GCGR conformations (Figure 2A, 2B, Supplementary Video 1). These correspond to a binding site between helices TM1 and TM2 (site-1), the extracellular portion of TM3/TM4 (site-2), an intracellular site formed by TM5, ICL3 and TM6 (site-3), and a densely packed site at the centre of TM6/TM7 (site-4). We note that for site-1, in the inactive conformation interacting residues were diffuse (Figure 2A), whereas in the active GCGR conformation they were restricted to the extracellular region (Figure 2B). The residues involved in formation of other sites were in broad agreement. There were small differences in the relative residence time order of the top four sites between the two receptor conformations. For the active GCGR conformation, cholesterol binding sites in proximity to TM5/TM6 (site-3 and site-4) were stabilised and site-2 interactions were destabilised when compared to the inactive conformation. These findings indicate that, whilst the location of key cholesterol binding sites is relatively insensitive to GCGR conformation, receptor activation induces subtle changes in cholesterol interactions that result in more prolonged residence times around TM5, ICL3, TM6 and TM7. Simulations performed using membrane bilayers with varying cholesterol content (Ansell et al., 2021) allowed us to estimate apparent dissociation constants for cholesterol at each site (Figure 2C): K_d_^app^ at site-1: 10.7 ± 0.3 %; site-2: 18.3 ± 0.4 %; site-3: 15.0 ± 0.8 %; and site-4: 6.6 ± 0.1 %.

**Figure 2:**
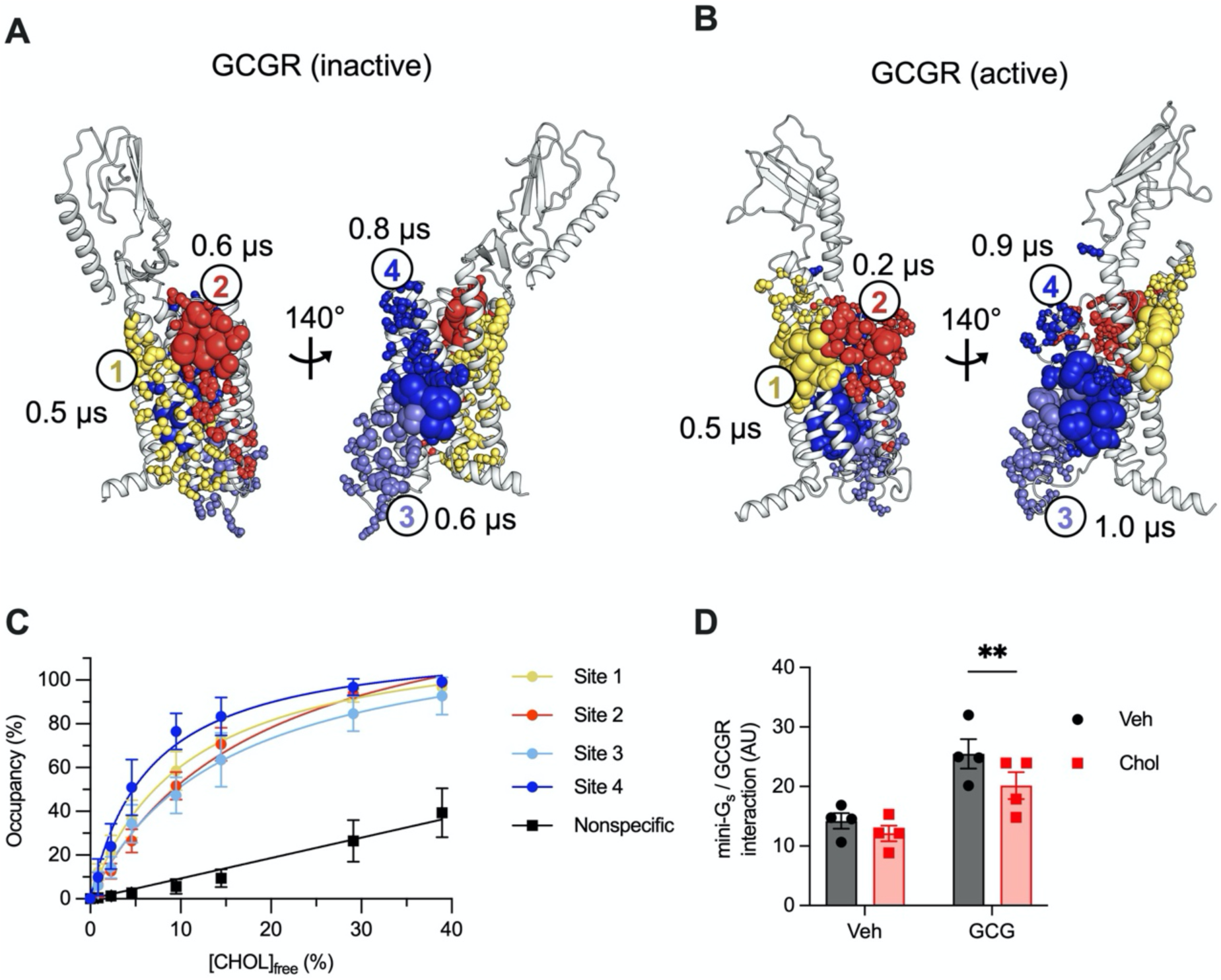
Predicted GCGR-cholesterol interactions. The top four ranked binding sites for cholesterol from coarse-grained MD simulations of the glucagon receptor (GCGR) in inactive (**A**) and active (**B**) conformations in plasma membrane-like bilayers containing 25% cholesterol. Each conformation was simulated for 10×10 μs. Distinct binding sites are coloured yellow (site-1), red (site-2), lilac (site-3) and blue (site-4). Residues comprising each site are shown as spheres scaled by per residue cholesterol residence times. The residence time for cholesterol binding to each site is indicated. Binding sites and associated residence times were calculated using PyLipID (Song et al., 2021). (**C**) Binding saturation curves for cholesterol binding to each site from equilibrium MD simulations (5x 5 μs at each % free cholesterol). Site % occupancy was calculated using PyLipID and plotted against the free cholesterol % (see methods) in binary bilayers composed of POPC and cholesterol. (**D**) Normalised AUC for luminescent signal indicating interaction between GCGR-SmBiT and mini-G_s_-LgBiT in HEK293T cells treated with 100 nM glucagon or vehicle for 30 min, *n*=4, compared by 2-way repeated measures ANOVA with Sidak’s test. ** p<0.01. Data are shown as mean ± SEM, with individual experimental replicates in Fig 2D.

Site-3 overlaps with the experimentally verified G protein binding site for GCGR and other class B GPCRs (Liang et al., 2017; Qiao et al., 2020). Moreover, site-4 corresponds to TM6, which tilts outwards during receptor activation to accommodate G protein binding. In view of the inverse correlation between membrane cholesterol content and cAMP signalling (Figure 1C), we hypothesised that cholesterol could act as a negative allosteric regulator at sites-3/4 (Abiko et al., 2021) by reducing the capacity of the receptor to interact with Gα_s._ This possibility was supported by studies measuring recruitment to the receptor of mini-G_s_, a conformational biosensor for Gα_s_-favouring active GPCR conformations (Nehme et al., 2017), in which cholesterol loading reduced glucagon-stimulated GCGR-mini-G_s_ interactions (Figure 2D).

### Increasing hepatic cholesterol in mice decreases glucagon sensitivity

Next we manipulated hepatic cholesterol in adult mice using isocaloric chow high or low in cholesterol, with or without simvastatin. While these diets had no effect on body weight or hepatic triglyceride content (Figures 3A, 3B), the cholesterol-enriched diet caused a dramatic increase in hepatic cholesterol after 1 week, which was partially abrogated by the inclusion of simvastatin (Figure 3C). Following an intra-peritoneal glucagon/pyruvate challenge test, we observed that the expected glucagon-induced peak in glycaemia was absent in mice that had been fed the high cholesterol diet, but partly restored in mice fed the combined cholesterol and simvastatin diet (Figure 3D). There was also an inverse association between hepatic cholesterol and the glucose excursion conferred by addition of glucagon (R^2^= 0.14; p = 0.02; Figure 3E).

**Figure 3:**
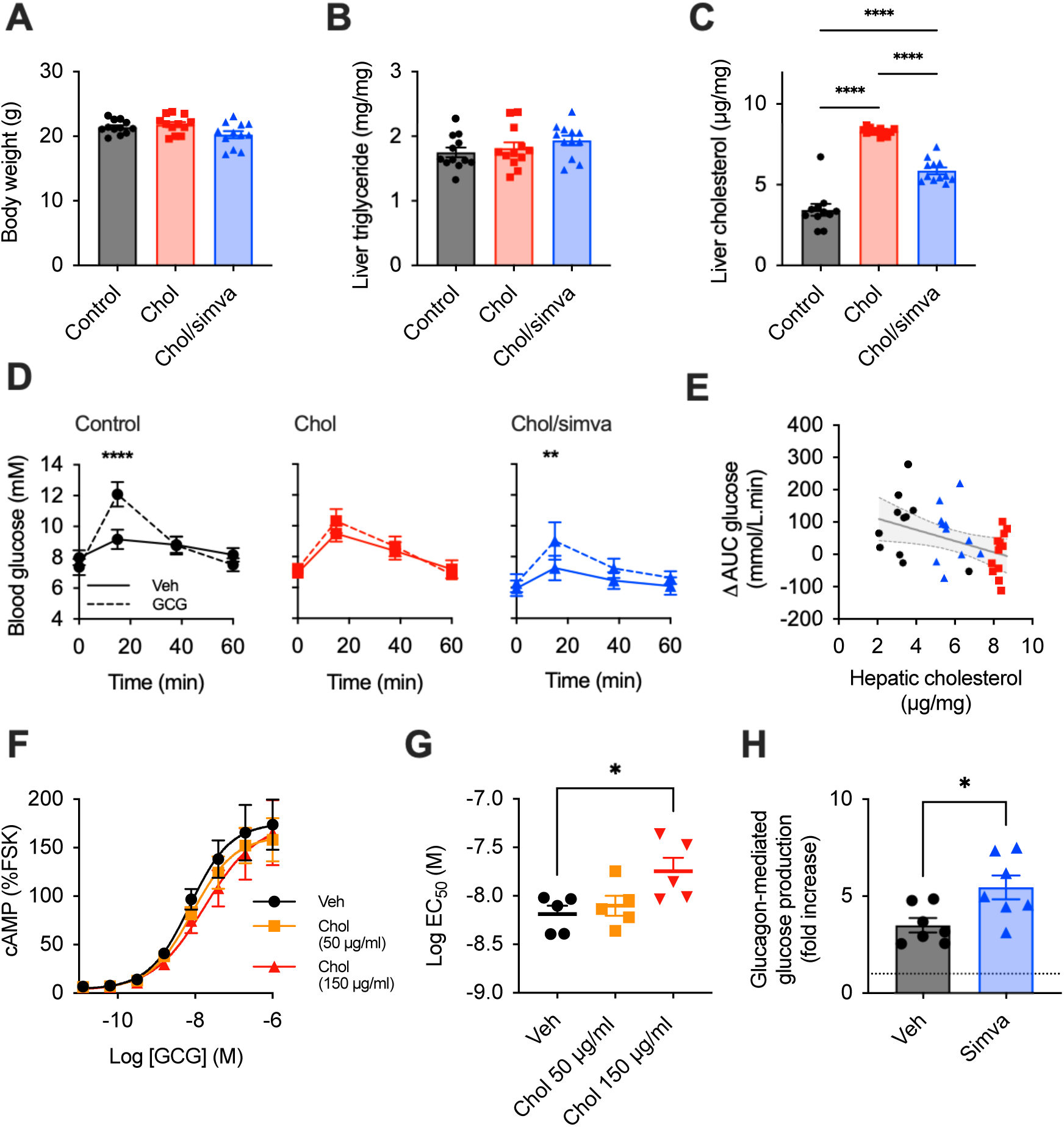
Increase in hepatic cholesterol in mice decreases responsiveness to glucagon. (**A**) body weight, (**B**) hepatic triglyceride, (**C**) hepatic cholesterol, (**D**) glucagon challenge test and (**E**) association between change in AUC during glucagon challenge test and hepatic cholesterol in mice fed different diets as indicated by colour code; the linear regression line ± 95% confidence intervals is shown; n=12 in each group. cAMP dose response curves (**F**) and log EC_50_ (**G**) in primary mouse hepatocytes pre-treated with cholesterol, then stimulated with glucagon; n=5, 4-parameter fit of pooled data shown; (**H**) glucose production in primary mouse hepatocytes pre-treated with simvastatin or vehicle overnight, then stimulated with glucagon for 24 hours, expressed as fold change over no-glucagon control stimulation, n=7. Data are represented as mean ± SEM with individual replicates shown in some cases; statistical analysis performed with one-way ANOVA and Sidak’s post-hoc test (A, B, C, G), two-way ANOVA and Sidak’s post-hoc test (D), or unpaired t-test (H).

*Ex vivo*, loading of wild-type primary murine hepatocytes with 150 µg/ml cholesterol led to a significant reduction in cAMP potency for glucagon, while 50 µg/ml had a more modest effect (Figure 3F, 3G). Prior treatment of primary hepatocytes overnight with simvastatin increased glucose output in response to glucagon stimulation (Figure 3H).

## Discussion

In this study we have demonstrated an inverse relationship between hepatocyte cholesterol and glucagon responsiveness, as measured *in vitro* by glucagon-stimulated cAMP and glucose production, and *in vivo* via the hyperglycaemic response to exogenous glucagon administration. Further, we have identified probable cholesterol binding sites on GCGR that could mediate these effects.

To our knowledge this is the first time the effect of cholesterol on GCGR function has been reported. Increasing membrane cholesterol enhances the function of some GPCRs, e.g. the α_1A_-adrenergic receptor, and diminishes that of others, e.g. the cannabinoid receptor 1 (CB1) and β_1_-adrenergic receptor (Abiko *et al*., 2021; Guixà-González et al., 2017). Interestingly, we previously showed that cholesterol depletion in pancreatic beta cells led to reductions in both cAMP signalling efficacy and ligand-induced endocytosis of the closely related GLP-1R (Buenaventura *et al*., 2019). This partly contrasts with our current GCGR results, in which cholesterol depletion reduced uptake of fluoresecent glucagon but increased cAMP signalling. The discrepancy may reflect inherent differences in the effect of cholesterol on the two receptors, though it is worth noting that GPCR function is also modulated by concomitantly-interacting membrane proteins, e.g. RAMP2 for GCGR (McGlone *et al*., 2021), and other membrane constituents that differ depending on cell type, which could in turn impact the role played by cholesterol. Other class B GPCRs for which the impact of cholesterol manipulation have been studied are summarised in Supplementary Table 1, with GCGR being somewhat unique within this class in showing enhanced cAMP signalling after cholesterol depletion.

There are various mechanisms by which cholesterol and other lipids may alter the stability, ligand binding properties, and thus function of GPCRs (Reiter *et al*., 2017): direct competition with agonist binding at the orthosteric site; directly binding at an allosteric site to modulate receptor conformation and dynamics (Abiko *et al*., 2021); indirectly via a change in local membrane composition and properties; or a combination of the above (Guixà-González *et al*., 2017). Recent work has demonstrated that GCGR function can be affected by endogenous allosteric modulators (Han et al., 2015; McGlone *et al*., 2021). GCGR has computationally predicted potential allosteric cholesterol binding sites, but the validity of these so-called “CRAC” and “CARC” consensus motifs in general has been questioned (Taghon et al., 2021). Our recently developed MD simulation method to evaluate GPCR-lipid interactions (Song *et al*., 2021) has allowed us to identify probable binding sites for cholesterol on the GCGR, with residence times which differ slightly depending on the receptor state. Site-4, which has the longest cholesterol residence time and lowest K_d_^app^, overlays with the observed binding of a negative allosteric regulator (Zhang et al., 2017) and antagonist (Jazayeri et al., 2016) in inactive GCGR structures. Cholesterol has been observed to bind to sites-1/2/3 in the structures of other G protein bound class B1 GPCRs, however site-4 is not observed in active-like conformations (Supplementary Table 2). Thus, we propose cholesterol binding to site-4 may be responsible for our observed decrease in glucagon responsiveness and mini-G_s_ binding by shifting the population ensemble toward the inactive conformation, similar to that proposed for other GPCRs (Casiraghi *et al*., 2016). Additionally it is possible that lipid synergy may further regulate receptor signalling. For example sites-3/4 are in proximity to previously observed PIP_2_ binding sites on GCGR (Ansell et al., 2020). Lipid interplay has been suggested to occur for anionic lipid binding to the ion channel Kir2.2 (Duncan et al., 2020a) and warrants further investigation in GPCRs.

An alternative mechanism for GCGR regulation may be receptor redistribution into distinct lipid nanodomains. Changes to the pool of accessible cholesterol within sphingomyelin enriched regions (Das et al., 2014), for example as a result of diet (Hermetet et al., 2019; Levental et al., 2016), may alter the combination of occupied cholesterol sites. Our observed mid-range K_d_^app^s are within the (patho)-physiological range of membrane cholesterol, rendering differential GCGR partitioning between lipid pools due to changes in cholesterol binding/unbinding a viable prospect.

Downstream of cAMP production, glucagon activates pathways that result in increased hepatic glucose production. Using dietary manipulation, we demonstrated that hepatic cholesterol content is inversely proportional to the hyperglycaemic response to a glucagon stimulus in mice. The effect observed was small, possibly due to compensatory mechanisms that make the investigation of glucagon sensitivity challenging *in vivo* (Janah *et al*., 2019). As glucagon exerts other effects on hepatic metabolism, including promoting amino acid and lipid catabolism, it would be valuable in the future to determine if these effects are similarly blunted. It is also unclear what the impact of cholesterol-mediated increases in GCGR internalisation could be, as suggested by our observations using fluorescent glucagon. GPCR internalisation is traditionally seen as a prequel to downregulation e.g. via lysosomal degradation, but sustained endosomal cAMP generation could alternatively lead to engagement with spatially constrained signalling networks not accessible to membrane-resident receptor (Manchanda et al., 2021).

Our study has limitations. Firstly, each of the experimental approaches we used to manipulate cholesterol have their own individual caveats. MβCD is likely to sequester additional lipids as well as cholesterol from the plasma membrane (Zidovetzki and Levitan, 2007), which could themselves influence receptor function (Ansell *et al*., 2020), although our study benefits from using lower MβCD concentrations than many others (see Supplementary Table 1). Inhibition of HMG-CoA reductase with statins reduces synthesis of not only cholesterol but also of intermediaries required for post-translational protein modifications including farnesylation and geranylgeranylation (Werner et al., 2002). The dietary changes we implemented in mice resulted in clear alterations to hepatic cholesterol content, but we cannot infer a plasma membrane-specific effect from this approach in isolation. Despite these caveats, we observed congruent results across different systems that support a role for cholesterol in the regulation of GCGR sensitivity. A second limitation of our study is that we are unable to state confidently whether manipulating the lipid environment of the GCGR influences its signalling properties primarily via a direct effect on receptor function, or by altering the spatial dynamics of GCGR relative to its intracellular effectors. These possibilities could be explored in follow up studies using, for example, membrane fractionation techniques, *in vitro* reconstituted systems, imaging approaches to co-visualise the receptor and its potential interactors, and solid-state NMR experiments at variable cholesterol concentrations.

There are several possible implications of our findings for human health and disease. Statin treatment is associated with an increased incidence of new-onset T2DM (Danaei et al., 2013; Wang et al., 2017) and progression of existing T2DM (Mansi et al., 2021), with the degree of low-density lipoprotein-cholesterol (LDL-C) reduction correlating with the likelihood of developing T2DM in these studies (Wang *et al*., 2017). Genetic polymorphisms of *HMGCR* and related genes that lead to reduced LDL-C also increase the probability of developing T2DM (Lotta et al., 2016; Swerdlow et al., 2015), whereas patients with monogenic familial hypercholesterolaemia have high levels of plasma and hepatic cholesterol but a reduced risk of incident T2DM (Besseling et al., 2015; Hoeg et al., 1984). Our data raise the intriguing possibility that these effects may occur in part via a reduction in hepatocyte membrane cholesterol increasing sensitivity to the hyperglycaemic effects of physiological glucagon.

Our data also potentially reconcile the observations that NAFLD is associated with both an increase in hepatic cholesterol (Min *et al*., 2012; Puri *et al*., 2007) and glucagon resistance (Albrechtsen et al., 2018; Gar *et al*., 2021). Contrasting with the possibility discussed above, i.e. that statin treatment might enhance glucagon sensitivity and thereby increase fasting blood glucose levels, glucagon resistance in NAFLD has paradoxically been proposed to drive impaired glucose tolerance and T2DM in a subset of patients. This may be via perturbation of the alpha cell-hepatocyte axis, whereby hepatocytes resistant to glucagon are less able to catabolise amino acids, with the resulting increases in plasma amino acid levels leading to hypersecretion of glucagon by alpha cells (Holst et al., 2017). In this context, hyperglucagonaemia is still capable of increasing glycaemia as glucagon resistance is incomplete. In view of these ostensibly opposite effects on glycaemia that could result from cholesterol-mediated glucagon resistance, further studies are needed to carefully examine which process dominates in different pathological states. Nevertheless, greater understanding of this relationship is likely to underpin new therapies for NAFLD (Younossi et al., 2021).

Antagonising glucagon signalling has long been proposed as a therapeutic strategy for T2DM (Cheng et al., 2020). Our data suggest that cholesterol binding site-4 could be targeted by small molecule allosteric modulators of GCGR activity, as has been achieved for other GPCRs (Lu et al., 2017). On the other hand, GCGR agonism is increasingly seen as a viable component of multi-incretin treatment for obesity and diabetes as, when combined with GLP-1R agonism, beneficial effects of glucagon (e.g. amelioration of hepatic steatosis and enhanced weight loss through increased energy expenditure) may be realised without unwanted hyperglycaemia (Ambery et al., 2018). Our work therefore suggests that it is worth evaluating whether lipid-modifying treatments can aid the effectiveness of therapeutic GCGR agonism in metabolic disease.

## Acknowledgements

ERM was supported by a UK Medical Research Council (MRC) Clinical Research Training Fellowship and an Early Career Grant from the Society of Endocrinology while working on this project. TBA is funded by the Wellcome Trust (102164/B/13/Z). AT and DC are funded by the MRC (MR/R010676/1 and MC-A654-5QB10, respectively). AT also acknowledges support from Diabetes UK and the European Federation for the Study of Diabetes. BJ is supported by the MRC (MR/R010676/1), IPPRF scheme, European Federation for the Study of Diabetes, Society for Endocrinology, British Society for Neuroendocrinology, and the UK National Institute for Health Research (NIHR) Imperial Biomedical Research Centre (BRC). TT is funded by the NIHR, the NIHR Imperial BRC and the J.P. Moulton Charitable Foundation. SRB is funded by the NIHR Imperial BRC. MS is supported by Wellcome Trust (208361/Z/17/Z and 220062/Z/20/Z) and the UK Biotechnology and Biological Sciences Research Council (BBSRC BB/R00126X/). The Department of Metabolism, Digestion and Reproduction at Imperial College London, UK, is funded by grants from the MRC, the BBSRC and is supported by the NIHR Imperial BRC. The views expressed are those of the authors and not necessarily those of the abovementioned funders, the UK National Health Service (NHS), the NIHR, or the UK Department of Health.

## Author contributions

Conceptualization: ERM, BJ, TT, SRB; Methodology: BJ, AT, WS, TBA, MS; Investigation: ERM, BJ, CD, TBA; Writing, reviewing and editing: all authors.

## Declaration of Interest

The authors declare no competing interests

## Materials and methods

### Cell culture and primary mouse hepatocyte isolation

Huh7-GCGR cells (McGlone *et al*., 2021) were cultured at 37°C in 5% CO_2_ in DMEM supplemented with 10% FBS, 1% penicillin/streptomycin and 1% G418 (Thermo Fisher). HEK293T cells were cultured similarly but without G418. Hepatocytes from male adult C57Bl/6J mice were isolated using collagenase perfusion, as previously described (Woods et al., 2011). After washing and resuspension, primary hepatocytes were plated and cultured at 37°C in 5% CO_2_ in Medium 199 supplemented with 1% penicillin/streptomycin, 1% BSA, 2% Ultroser G, 100 nM T3, 100 nM dexamethasone and 100 nM insulin (all Thermo Fisher). After 5 hours, medium was changed to M199 with 1% penicillin/streptomycin, 100 nM dexamethasone and 10 nM insulin for serum starvation.

### Peptides

Glucagon and a fluorescent glucagon analogue, “FITC-GCG”, which conforms to the amino acid sequence of glucagon(1-29) with a C-terminal extension Gly30,31Lys32-FITC (fluorescein isothiocyanate) (McGlone *et al*., 2021), were obtained from Wuxi Apptec.

### Cholesterol-modulating treatments

Cells in suspension or plated were treated with cholesterol-saturated methyl-β-cyclodextrin (referred to as “cholesterol” hereafter when in the context of cellular treatments; at 50 µg/ml unless otherwise stated), or cholesterol-depleted methyl-β-cyclodextrin (MβCD; varying concentrations, Sigma Aldrich), in Hanks’ buffered saline solution (HBSS), whilst being gently agitated or rocked for 30 minutes. Plated cells were treated with simvastatin 10 µM and/or mevalonate 50 µM (Sigma-Aldrich) for 16 hours in serum-free medium. All treatments were thoroughly washed off prior to experiments.

### cAMP accumulation assay

After cholesterol-modulating treatments as described above, Huh7-GCGR cells in 96-well plates were stimulated with indicated concentration of agonist in serum-free medium for 10 min at 37°C. Where indicated, 10 µM forskolin was used as a positive control, and results were normalised to the forskolin response. Where indicated, 10 ng/ml pertussis toxin (PTX; Sigma-Aldrich) was applied overnight in advance of the assay to inhibit Gα_i_. Primary mouse hepatocytes in suspension were stimulated in serum-free DMEM with phosphodiesterase inhibitors (100 µM 3-isobutyl-1-methylxanthine; IBMX) for 10 minutes at 37°C. After stimulation cells were lysed and cAMP was assayed by immunoassay (Cisbio HTRF cAMP Dynamic 2).

### Fluorescent glucagon uptake assay

After cholesterol-modulating treatments as described above, Huh7-GCGR cells in 96-well black, clear bottom plates were stimulated with FITC-GCG for 10 minutes at 37°C in serum-free medium. Unbound agonist was removed by washing three times with HBSS, followed by fixation with 2% paraformaldehyde (PFA) for 10 minutes at room temperature. High content imaging was performed using an automated Nikon Ti2 widefield microscope with a 0.75 numerical aperture 20X air objective, with 9 fields-of-view acquired per well, and internalised agonist was quantified from epifluorescence images as previously described (Pickford et al., 2021).

### Mini-G_s_ recruitment assay

HEK293T cells were co-transfected using Lipofectamine 2000 with human GCGR bearing an N-terminal SNAP-tag and C-terminal SmBiT tag, plus LgBiT-tagged truncated Gα_s_ subunit (mini-G_s_-LgBiT, a gift from Prof Nevin Lambert, Medical College of Georgia), as previously described (Jones et al., 2020). At 36 hours, an equal number of detached cells were treated in suspension with cholesterol for 30 min. After further washing, cells were resuspended in HBSS containing Furimazine (1:50 dilution; Promega) and dispensed into 96-well half-area white plates. After reading baseline luminescence over 8 min, 100 nM glucagon or vehicle (HBSS) was added and signal serially recorded for 30 min. Signal was normalised to the average baseline across all conditions within each experiment, and integrated area-under-curve was calculated.

### Molecular dynamics simulations and analysis

Structures of the GCGR in inactive and active conformations were derived from the Protein Data Bank (PDB ID: inactive 5XEZ, active 6LMK) (Qiao *et al*., 2020; Zhang *et al*., 2017). Additional components (including bound glucagon in 6LMK) were removed and missing loops and/or incomplete residues were modelled using Modeller (Ansell *et al*., 2020; Fiser and Sali, 2003). The GCGR was coarse-grained using *martinize*.*py* (de Jong et al., 2013) and embedded in an asymmetric lipid bilayer membrane comprising POPC (20%), DOPC (20%), POPE (5%), DOPE (5%), sphingomyelin (15%), GM3 (10%) and cholesterol (25%) in the upper leaflet and POPC (5%), DOPC (5%), POPE (20%), DOPE (20%), POPS (8%), DOPS (7%), PIP_2_ (10%) and cholesterol (25%) in the lower leaflet using *insane*.*py* (Wassenaar et al., 2015). The MARTINI2.2 coarse grained (CG) forcefield (Marrink et al., 2007) was used to describe all components and cholesterol was modelled with the virtual site descriptor (Melo et al., 2015). The ElNeDyn elastic network was applied to each protein with a cut-off of 0.9 nm and a spring force constant of 500 kJ mol^-1^ nm^-2^ (Periole et al., 2009). Each system was solvated using MARTINI water (Marrink *et al*., 2007) and ∼0.15 M NaCl before independent minimization and equilibration steps. Simulations were run identically to those described previously (Ansell *et al*., 2020) for 10x 10 μs of each GCGR conformation. The GROMACS 5.14 and 2019 simulation packages were used to perform simulations (www.gromacs.org). Cholesterol interactions with GCGR were analysed using PyLipID (Song *et al*., 2021) with a 0.55/0.8 nm double cut-off to define lipid contact. Binding sites and kinetic analysis (i.e. derived residence times) were calculated by PyLipID as described in (Song *et al*., 2021).

### Computational binding saturation curves

Binding saturation curves and site affinities were calculated in accordance with a recently published method (Ansell *et al*., 2021). GCGR (PDB ID: active 6LMK) was simulated in binary POPC:cholesterol bilayers of increasing cholesterol content (1%, 2.5%, 5%, 10%, 15%, 30%, 40%). Each system was simulated for 5x 5 μs with CG simulation parameters as described above. The concentration of free cholesterol (%) was defined as the mean number of unbound cholesterol (> 0.8 nm from the GCGR transmembrane domain) divided by the total number of lipids in the bilayer. Site occupancies correspond to the mean occupancy of 6 residues at each site, as obtained by PyLipID. Apparent dissociation constants (K_d_^app^s) were calculated via fitting to equation 1 using Prism 9.2.0 for MacOS (Graphpad Software).

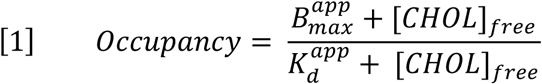

### Hepatocyte glucose output assay

Primary hepatocytes were serum-starved overnight with the addition of simvastatin 10 µM or vehicle (DMSO). After a thorough wash to remove any phenol-red containing media, glucose production media was added to the wells (phenol-red free DMEM with 1 mM sodium pyruvate (Gibco) and 20 mM sodium lactate (Sigma). Baseline samples were taken before addition of glucagon 100 nM or vehicle. The plate was incubated at 37°C for 24 hours, after which media was removed and stored at −20°C. Cellular protein was extracted and quantified using a BCA assay (Thermo Fisher). Glucose in media was assayed using a glucose oxidase assay (Randox), normalised to protein content of the well, and expressed as fold change of glucose production in the absence of glucagon.

### Animals and dietary manipulation

Experiments were performed in accordance with the UK Animals (Scientific Procedures) Act 1986 and approved by the Animal Welfare and Ethical Review Board at Imperial College London. C57BL/6J male mice (Charles River) were group-housed in a pathogen-free facility at controlled temperature (22°C) with a 12-hour light dark cycle. Access to food was ad libitum, except prior to fasting studies, and mice always had free access to water. Mice were fed standard chow (SDS Rm3) during acclimatisation. Specialist chows were based on a standard Clinton/Cybulsky rodent diet (10% kcal from fat and 70% kcal from carbohydrate) and were identical in constitution and calorie content, with the exception of 0% or 0.5% added cholesterol, or 0.5% cholesterol and 120 mg simvastatin per kg (Research Diets). Mice were fed specialist diets for at least 7 days before glucagon challenge test and 12 days before the cull.

### Glucagon challenge test

Glucagon challenge tests was performed in the light phase, in mice fasted for 5 hours (Gelling et al., 2003; Rossi et al., 2018). Tail vein blood glucose was measured using a handheld glucometer (Nexus, GlucoRx) before and at intervals following intraperitoneal injection with sodium pyruvate 2 g/kg (Sigma) as a gluconeogenic substrate ± glucagon 10 nmol/kg. Each mouse underwent the test with and without glucagon, in a random order, with an intervening washout period of 3 days. At the end of the study mice were culled using decapitation following a 5-hour fast. The liver was harvested, snap frozen in liquid nitrogen and stored at −80°C.

### Lipid extraction

Lipids were extracted from liver tissue by homogenization in ethanol (0.03x w/v) (Woods et al., 2017). Lipids were extracted from cells by agitation in butanol, before evaporation and resuspension in methanol.

### Cholesterol and triglyceride assays

Triglyceride content of samples was measured using a GPO-PAP Triglyceride assay (Randox Laboratories Ltd) and cholesterol using Amplex Red Cholesterol Assay Kit (ThermoFisher Scientific).

### Statistical analyses

All statistical analysis of experimental data was performed using Prism 9.0. Concentration-response curves were generated by 3-parameter logistic fitting or using a biphasic “bell-shaped” fit. For cAMP, E_max_ and log_10_-transformed EC_50_ were derived for each repeat and then compared using t-tests or one-way ANOVA, with matched analyses performed where permitted by experimental design, and multiple comparison tests as indicated in the figure legends. For experiments using PTX, the Gα_i_ component of the biphasic response was calculated by subtracting response in the presence of PTX (i.e. the Gα_s_-specific response) from total response. For cellular treatments, as a combined measure of agonism, E_max_/EC_50_ was calculated and normalised to vehicle control; log_10_-transformed values were then used for simple linear regression analysis, with calculation of goodness of fit. p<0.05 was considered statistically significant.

---

**Supplementary Table 1:** Class B GPCRs for which the impact of membrane cholesterol modulation has been reported.

| Class B GPCR | Cell model | Manipulation and effect |
| --- | --- | --- |
| Calcitonin gene-related peptide receptor (CGRPR) | Guinea pig gall bladder (Jennings et al., 1999) smooth muscle | Cholesterol-saturated MβCD: ↓ CGRP-induced action potentials |
| Glucagon-like peptide receptor 1 (GLP-1R) | INS-1 cells (Buenaventura *et al.*, 2019) | 10 mM MβCD: ↓ exendin-4-induced cAMP efficacy and endocytosis |
| | HEK293 cells (Lucey et al., 2021) | 10 mM MβCD: ↓ exendin-4-C16-induced cAMP potency and endocytosis |
| Glucagon-like peptide 2 receptor (GLP-2R) | DLD-1 and BHK cells (Estall et al., 2004) | 10 mM MβCD: ↓ GLP-2-induced endocytosis, no effect on cAMP |
| Glucagon receptor (GCGR) | Huh7 cells and mouse hepatocytes (this work) | 1 – 10 mM MβCD, statin treatment: ↑ glucagon-induced cAMP, ↓ endocytosis; opposite effect from cholesterol loading |
| Pituitary adenylate cyclase-activating polypeptide type 1 receptor (PAC<sub>1</sub>R) | PC12 (Emery et al., 2015) | 10 mM MβCD: ↓ PACAP-38 cAMP efficacy |
| Parathyroid hormone type 1 receptor (PTHr1) | HEK293 cells (Tovey and Taylor, 2013) | 2% MβCD: no effect on PTH-induced augmentation of carbachol-induced Ca<sup>2+</sup> transients |
| Vasoactive intestinal polypeptide receptor 2 (VPAC<sub>2</sub>R) | Mouse gastric smooth muscle cells (Mahavadi et al., 2013) | 10 mM MβCD: ↓ VIP-induced endocytosis |

**Supplementary Table 2:** Class B GPCRs with deposited cholesterol binding sites on RCSB Protein Data Bank (https://www.rcsb.org; accessed 13^th^ October 2021)

| Class B GPCR | Structures with modelled cholesterol | Structures with cholesterol bound at site-1, -2 or -3 |
| --- | --- | --- |
| Parathyroid hormone type 1 receptor (PTH1R) | 6NBF, 6NBI, 6NBH (Zhao *et al.*, 2019) | 6NBF (site-2), 6NBH (site-2, site-3) |
| Glucagon-like peptide receptor 1 (GLP-1R) | 7DUQ, 7E14 (Cong *et al.*, 2021) | 7DUQ (site-1, site-3), 7E14 (site-3) |
| Corticotrophin-releasing factor 1 and 2 receptors (CRF1R and CRF2R) | 6PB0, 6PB1 (Ma *et al.*, 2020) | 6PB0 (site-2), 6PB1 (site-1, site-2) |
| Glucose-dependent insulinotropic polypeptide receptor (GIPR) | 7DTY (Zhao *et al.*, 2021) | 7DTY (site-2, site-3) |
| Growth hormone releasing hormone receptor (GHRHR) | 7CZ5 (Zhou *et al.*, 2020) | N/A |
| Vasoactive intestinal polypeptide receptor 1 (VIP1R) | 6VN7 (Duan et al., 2020) | 6VN7 (site-2, site-3) |

## Notes

### Competing Interest Statement

The authors have declared no competing interest.

